# Cryo-EM and X-ray structures of an anti-MHC-I monoclonal antibody/H2-K^b^ complex reveal the basis of conformational dependence and allelic specificity

**DOI:** 10.64898/2026.09.11.750949

**Authors:** Jiansheng Jiang, Kannan Natarajan, Reanne Towler, Lisa F. Boyd, Allison B. Lupatkin, Rick K. Huang, David H. Margulies

**Affiliations:** Molecular Biology Section, Laboratory of Immune System Biology, National Institute of Allergy and Infectious Diseases, National Institutes of Health, Bethesda, Maryland, USA; Laboratory of Cell Biology, Center for Cancer Research, National Cancer Institute, National Institutes of Health, Bethesda, Maryland, USA

**Keywords:** antibody, major histocompatibility complex (MHC), surface plasmon resonance (SPR), cryo-electron microscopy (cryo-EM), X-ray crystallography

## Abstract

Monoclonal antibodies (mAbs) directed against MHC class I molecules (MHC-I) have been invaluable for a host of functions including tissue typing for transplantation, exploration of immune activation, and probing molecular structure and dynamics. We previously studied a set of murine mAbs directed against MHC-I to characterize their domain and allele specificity and demonstrated the consistency of structural identification of epitopes with prior mapping studies. Here, we report cryo-EM and X-ray crystal structures of a novel mAb (AF6-88.5) in complex with the mouse MHC-I molecule H2-K^b^. We characterize the conformational epitope that is sensitive to polymorphic amino acid residues in the MHC-I extracellular domains. The structure of the mAb/MHC-I complex reveals contacts of the mAb to residues of the α2, α3, and β2m domains and comparison to the unliganded mAb shows slight adjustments of the mAb CDRH3 and CDRL3 loops to accommodate the epitope surface. The structurally identified site overlaps with that seen by some coreceptors and immunoevasins. Amino acid sequence alignments and site-directed mutagenesis of H2-K^b^, H2-D^b^, and β_2_m explain the allelic specificity. These results emphasize the remarkable ability of antibodies to bind conformationally defined epitopes with exquisite discrimination and the contribution of MHC-I polymorphisms to antigenicity.

## Introduction

Monoclonal antibodies (mAbs) directed against MHC-I molecules are invaluable diagnostic, therapeutic, and research tools both clinically and experimentally. Since the advent of hybridoma technology, many MHC-I allele-specific mAbs have been developed and are used widely to detect, quantify, or block cell specific surface MHC-I molecules in various experimental and therapeutic settings. Several such mAbs can discriminate between closely related amino acid sequences and thus offer a rich trove of proteins for investigating the molecular basis of biological specificity and its manipulation for clinical benefit. Antibodies provide a remarkable example of the adaptability of a fundamental protein structural scaffold to permit sufficient variability to bind a host of distinct antigenic structures with specificity and fidelity (1). The development of monoclonal antibody technologies and the subsequent application of a variety of recombinant display approaches have fostered the ability to identify specific reagents that interact with almost any biochemical structure, often with great specificity (2, 3). An ever-expanding database of experimentally determined structures of antibody/protein antigen complexes now includes about 7500 examples (https://sabdab.opig.stats.ox.ac.uk/statistics (4)), but the ability to computationally predict antibody epitopes and to design antibodies with well-defined epitopic specificity remains a major technological challenge (5). Among protein antigens that play critical roles in a variety of recognition phenomena that regulate the immune response are the cell surface molecules encoded by the major histocompatibility complex (MHC), designated HLA (human leukocyte antigen) class I or class II in the human or H2 or Ia in the mouse. MHC-I molecules are of particular interest because of their remarkable polymorphism (more than 30,000 alleles have been identified in the human (6)). MHC-I molecules are ubiquitously expressed cell-surface trimers consisting of a polymorphic HLA or H2 heavy chain of about 45 kDa, a β_2_-microglobulin (β_2_m) light chain of about 12 kDa, and a bound repertoire of 8-12-mer peptides representative of the cell of biosynthesis. Recombinant MHC-I molecules can be prepared with individual peptides (7). MHC-I molecules function at the cell surface to interact with receptors on natural killer (NK), various myeloid, and T cells. We have recently explored structures of several previously characterized anti-MHC-I mAbs by X-ray crystallography and cryo-electron microscopy (cryo-EM) and have rationalized their binding specificities and functional effects based on the structural identification of their epitopes (8–10). Several of these anti-MHC-I antibodies exert profound immunological effects in mice or humanized mouse disease models (9–12). Although computational approaches to define epitopic sites of protein antigens identified by specific antibodies have shown considerable improvement in recent years (13–15), fine details of mAb/protein interactions are predicted with low accuracy, demanding further augmentation of the database of experimentally defined structures of mAb/protein antigen complexes.

Among a host of anti-MHC-I mAbs that have been characterized by various epitope-mapping techniques, those with precise allele-specific reactivities are of particular interest. One such antibody, AF6-88.5, directed against the mouse allelomorph, H2-K^b^, used widely in serological and functional studies, has been reported to be highly selective for H2-K^b^ heavy chains complexed with murine β_2_m (mβ_2_m), conformationally sensitive, and peptide dependent but not peptide specific (16–20). It also plays a critical role in mouse antibody models for transfusion associated acute lung injury (TRALI) where it functions synergistically with another anti-H2 mAb (34-1-2S) to promote disease in C57BL/6 mice (21). Here, we present binding affinity data, a cryo-EM structure of the Fab AF6-88.5/H2-K^b^ complex and X-ray structures of the AF6-88.5 mAb, both alone and in complex with H2-K^b^. These results, verified by site-directed mutagenesis, explicitly define the MHC-I epitope seen by the mAb and also explain its conformational dependence and allelic specificity. The structure of the complex provides a topological explanation for the contribution of AF6-88.5 to the mouse model of TRALI and its proposed molecular mechanism.

## Results

### Binding specificity of AF6-88.5

The anti-H2-K^b^ mAb AF6-88.5 (20) is specific for peptide-bound (18) mβ_2_m-complexed (16, 22) molecules, and previous mutational studies suggested that the epitope was influenced by amino acid residues of each of the three extracellular domains of H2-K^b^ (16), an intriguingly broad epitope considering the single allelic specificity of this mAb. We examined the MHC-binding specificity of this mAb using surface plasmon resonance (SPR) and measured the interaction with a selection of recombinant murine and human MHC-I molecules, prepared with either the mouse or human β_2_m light chain (Fig. 1, *A* and *B*). As shown in Fig. 1*A*, H2-K^b^ complexed with mβ_2_m and peptide (SIINFEKL) binds strongly, whereas complexes containing hβ_2_m bind poorly. There was minimal binding of H2-D^b^/mβ_2_m or of H2-L^d^/mβ_2_m complexes, and minimal binding of HLA-A*02:01, H2-K^d^, or of mouse H2-D^b^/hβ_2_m, or of mβ_2_m or hβ_2_m alone (Fig. 1*A*). We evaluated the binding affinity (*K*_D_) of the AF6-88.5/H2-K^b^ interaction to be 0.391 ± 0.05 μM (Fig. 1*B*), a value consistent with that previously reported (21).

**Figure 1.**
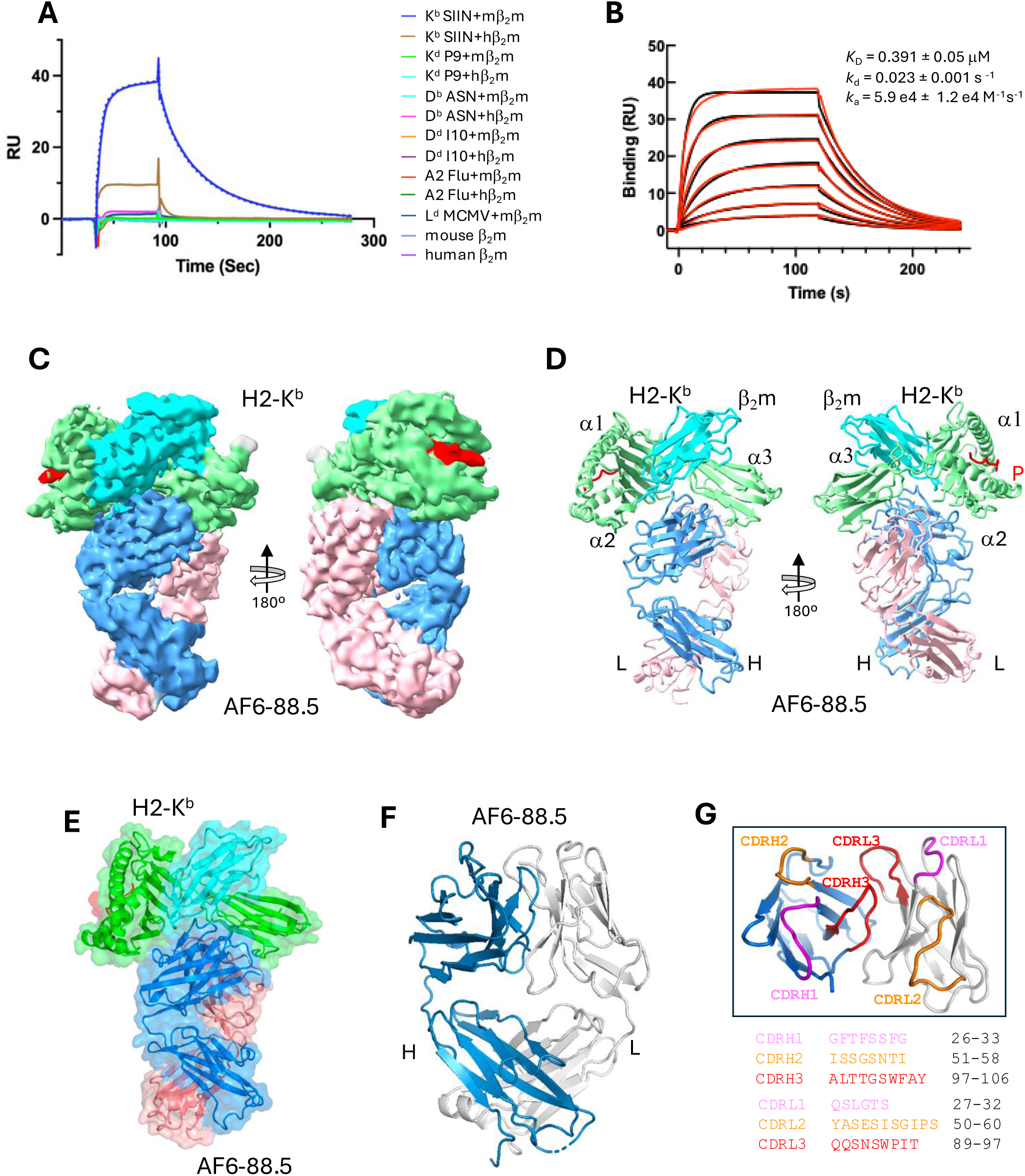
AF6-88 binds H2-K^b^ complexed with mβ_2_m and SIINFEKL. *A,* BIAcore CM-5 chip was coupled to mAb AF6-88.5 as described in **Experimental procedures**. Preparations of the indicated purified recombinant complexes were injected over the surface at the indicated point, and buffer washout was subsequently initiated at t=90 s. The blue and black dotted lines indicate the exposure of the surface to the same concentration of K^b^/mβ_2_m/SIINFEKL as the first and final cycles (representative set of binding curves, n=3). *B,* Affinity measurement based on kinetic binding by SPR. Binding was for 120 s before washout was initiated (n=3). *C,* Cryo-EM map of AF6-88.5+H2-K^b^ at resolution 3.09 Å (EMD-77500). *D,* Model of AF6-88.5+H2-K^b^ (PDB-ID: 36FZ) that fits the map. Lime color as α-chain and cyan color as β-chain of H2-K^b^ ; cornflower as heavy chain and pink as light-chain of AF6-88.5. *E,* X-ray crystal structure of AF6-88.5+H2-K^b^ (PDB-ID: 9Q0K) at resolution 3.44 Å. Green is α chain of H2-K^b^, cyan is β_2_m, and blue and salmon are heavy and light chain of AF6-88.5 respectively. *F,* X-ray crystal structure of Fab AF6- 88.5 (PDB-ID: 9Q0L) at resolution 2.60Å, surface representation. Blue is the heavy chain and gray is the light chain. *G,* CDR loops of AF6-88.5 are colored as pink for CDR1, orange for CDR2 and red for CDR3 respectively both for heavy and light chains.

### Cryo-EM Structure of the Fab-AF6-88.5/H2-K^b^ complex

To define the structural basis of the interaction of AF6-88.5 with H2-K^b^ we evaluated the complex by both cryo-EM and X-ray crystallography. Since the cryo-EM structure of the complex could be refined to somewhat higher resolution (3.09 Å) than the X-ray structure (3.44 Å), we discuss the cryo-EM structure first (Fig. 1, *C* and *D*) and then compare that with the X-ray crystal structure of the complex (Fig. 1*E*) and with that of the unliganded AF6-88.5 Fab (Fig. 1*F*). Details of cryo-EM data collection and data processing are presented in Table 1 and Fig. S1 and are described in **Experimental procedures**. We collected 4,356 movies (Fig. S1*A*) and used CryoSPARC^TM^ (23) for processing. Fig. S1*B* clearly shows the domains of AF6-88.5 and H2-K^b^ in the enlarged 2D class average (50x 2D classification). Application of the Multiple Low-Pass Filter (MLPF) protocol as described earlier for small Fab complexes (11) improved the map resolution to 3.09Å (Fig. S1*C* and *E*). Data collection statistics, refinement, and model quality for all three structural models are presented in Table 1 (cryo-EM) and Table 2 (X-ray). The overall structure of the complex reveals that the Fab is positioned with its complementarity determining region (CDR) loops (Fig. 1*G*) of both the heavy and light chains focused primarily on the membrane proximal α3 domain of H2-K^b^ with additional contacts to residues of the α2 domain and to the β_2_m subunit (Fig. 2 and Table 3). The Fab buries a total of 1300 Å^2^ of surface area of the H2-K^b^ H chain as well as 245 Å^2^ of β_2_m. Fig. 2*A* illustrates the footprints of AF6-88.5 on the H2-K^b^ surface, indicating where each CDR loop of the Fab recognizes the epitope on H2-K^b^ (Fig. 2*B, C,* and *D*). Inserts show the details of interactions of the CDR loops and their epitopes. The contact table (Table 3) reveals that AF6-88.5 CDRH1, 2, and 3 contact the H2-K^b^ H chain with contacts of CDRH1 to residues of the α3 domain (Q226, M228, E229, and L230) (Fig. 2*C*), of CDRH2 to α2 (beneath the peptide binding groove, residues R111, Y113, E128, T134) (Fig. 2*B*). AF6-88.5 CDRH2 has a broad interaction with two regions of β_2_m (T28/Q29; and S57/K58/D59) (Fig. 2*D* and Fig. 3*C*). The AF6-88.5 L chain also contacts H2-K^b^α3 (residues D212, Q218, G221, E222, E223, Y262 and P269 -- primarily through CDRL2) (Fig. 2C and Fig. 3B). The details of these interactions are shown in Fig. 2*C* and *D.* The H-2K^b^/β_2_m/peptide structure, as expected, is almost identical to a previously determined X-ray structure complexed with the same SIINFEKL peptide (PDB 3P9L), with RMSD of the H chain of 1.748 Å (for the first molecule in the asymmetric unit (AU) or 1.449 Å (for the second), with the differences primarily related to hinge angle flexibility between the α2 and α3 domains.

**Figure 2.**
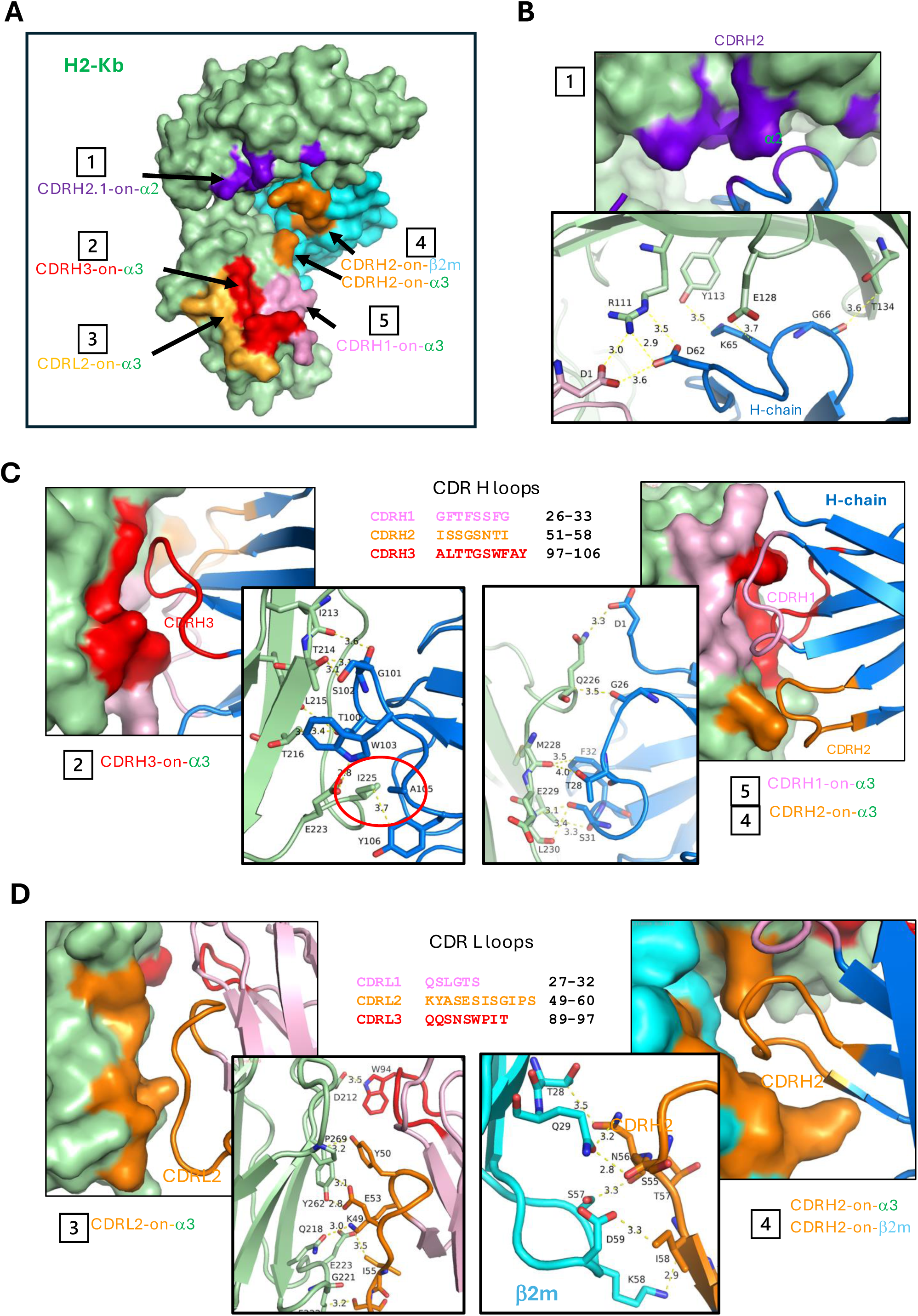
AF6 footprints on H2-K^b^ and epitopes. *A,* AF6-88.5 Fab footprints and interfaces on H2-K^b^ surface. (1) CDRH2 (D62, K65, G66) contacts with α2, (2). CDRH3 on α3, (3) CDRL2 on α3, (4) CDRH2 on both β2M and α3, (5) CDRH1 on α3. *B,* CDRH2 (D62, K65, G66) contacts with α2, the insert indicates the details of the interactions, as indicated in *A* (1). *C,* Three CDR loops of heavy chain of AF6-88.5 contact with α3 domain of H2-K^b^, inserts show details of interactions between H chain of AF6-88.5 and H2-K^b^, as indicated in *A* (2),(4),(5). Notice that I225 of α3 of H2-K^b^ interacts with the hydrophobic pocket formed by F27 and F32 of CDRH1, and A105 and Y106 of CDRH3. *D,* CDRL2 and CDRH2 contacts with α3 of H2-K^b^ respectively, inserts indicate details of interactions, as shown in *A* (3), (4).

**Figure 3.**
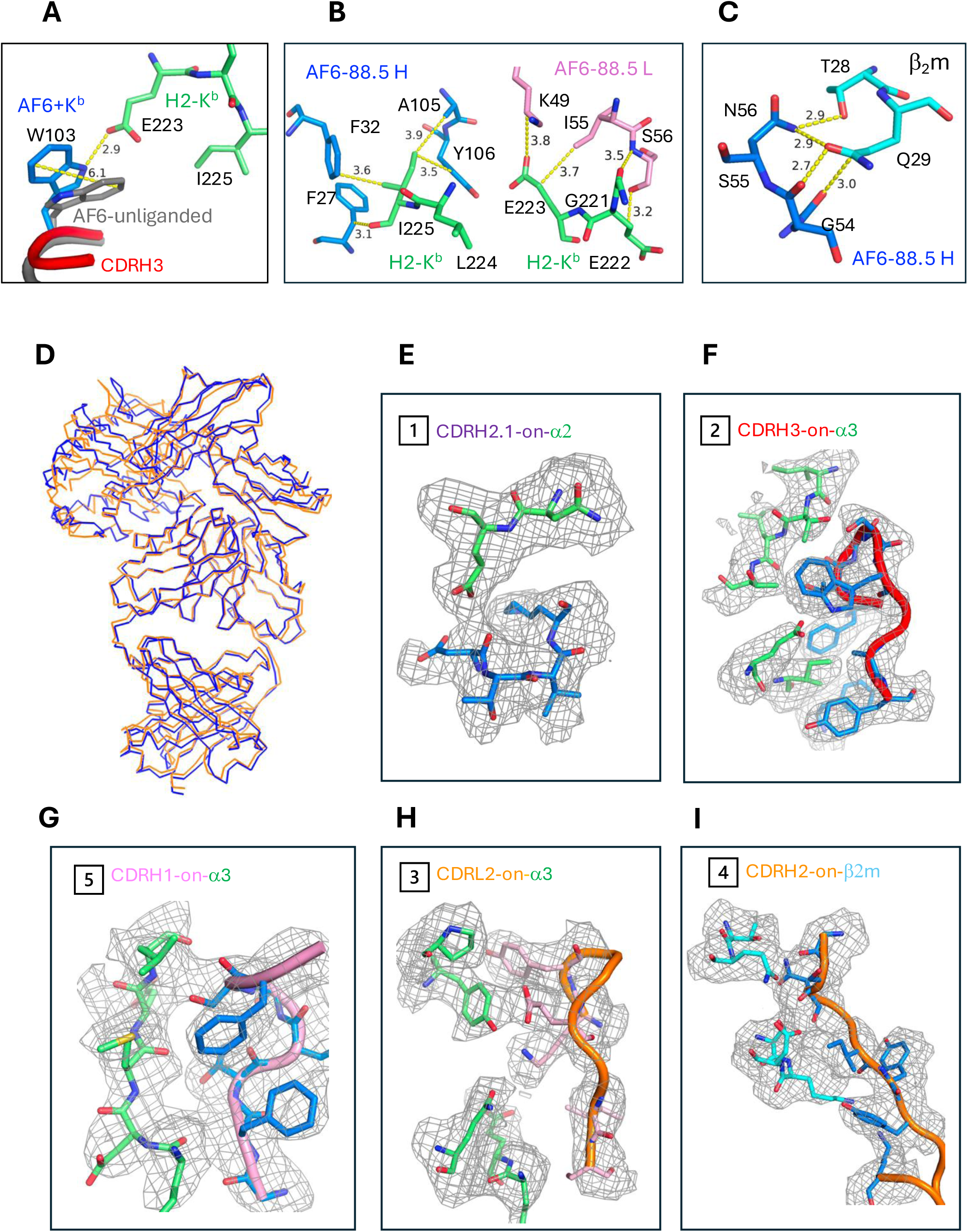
Superpositions reveal similarlity of cryo-EM and X-ray structures and minor changes in bound and free mAb loops. *A*, Comparison AF6-88+H2-K^b^ (blue) and AF6-88 unliganded (gray). The sidechain of W103 flipped leads to formation of a hydrogen bond between NC2 and NE2 of E223. *B*, AF6-88 recognizes the epitope of loop G221-I225 of H2-K^b^. I225 plays a vital role interacting with F27, F32, and Y106. *C*, G54-N56 of the CDRH1 loop recognize the epitopes of T28 and Q29 of β_2_M. *D,* Comparison cryo-Em and X-ray crystal structure. Superposed Cryo-EM (36FZ, blue) and X-ray structures (9Q0K, orange), represented as ribbons. RMSD=1.39Å. *E-I*. Cryo-EM (36FZ) density maps at the binding interface: *E* (1) CDR2.1 on α2, *F*.(2) CDRH3 on α3, *G*.(5) CDRH1 on α3, *H*.(3) CDRL2 on α3, *I.*(4) CDRH2 on β2M. The density is contoured at 1.5 σ.

**Table 1.** Cryo-EM data collection, refinement and validation statistics.

|  | **AF6-88Fab+H2-K<sup>b</sup>** |
| --- | --- |
| **EMDB ID** | EMD-77500 |
| **PDB-ID** | 36FZ |
| **Data collection and processing** |  |
| Magnification | 105,000 |
| Voltage (kV) | 300 |
| Electron exposure (e<sup>-</sup>/Å) | 54.2 |
| Defocus range (mm) | -0.7 to -2.0 |
| Pixel size (Å/pixel) | 0.83 (binned) |
| Raw micrographs (no.) | 4,356 |
| Extract particles (no.) | 2,326,183 |
| Refined particles (no.) | 372,395 |
| Particles for final map (no.) | 372,395 |
| Symmetry imposed | C1 |
| Map resolution (Å) | 3.09 |
| FSC threshold | 0.143 |
| **Refinement** |  |
| * Map sharpening B factor (Å<sup>2</sup>) | 0 |
| **Model composition** |  |
| Atoms | 6,241 |
| Residues | 808 |
| Ligands | 0 |
| **Overall B-factor (Å<sup>2</sup>)** |  |
| Protein (min/max/mean) | 90.9/298.1/170.2 |
| **R.m.s. deviations** |  |
| bond length (Å) | 0.002 |
| bond angle (°) | 0.49 |
| **CC (mask/volume/peaks)** | 0.73/0.73/0.62 |
| **Validation** |  |
| MolProbity score | 2.06 |
| Clash score | 6.85 |
| Ramachandran outliers (%) | 0 |

**Table 2.** X-ray Data collection and refinement statistics.

|  | **Fab AF6-88.5+H2-K<sup>b</sup>(PDB: 9Q0K)** | **AF6-88.5 Fab (PDB: 9Q0L)** |
| --- | --- | --- |
| Resolution range | 43.31 - 3.44 (3.71 - 3.44) | 40.66 - 2.6 (2.67 - 2.6) |
| Space group | P 21 21 21 | P 1 21 1 |
| Unit cell | 54.18 76.26 221.31 90 90 90 | 72.28 134.34 97.50 90 95.87 90 |
| Total reflections | 152090 (29808) | 427386 (28996) |
| Unique reflections | 12677 (2420) | 140763 (9725) |
| Multiplicity | 6.5 (6.6) | 3.0 (3.0) |
| Completeness (%) | 94.22 (90.37) | 96.92 (97.53) |
| Mean I/sigma(I) | 4.06 (1.23) | 2.83 (0.71) |
| Wilson B-factor | 81.82 | 50.12 |
| R-merge | 0.846 (2.715) | 0.4241 (2.027) |
| R-meas | 0.9202 (2.948) | 0.5156 (2.476) |
| R-pim | 0.358 (1.137) | 0.289 (1.399) |
| CC1/2 | 0.931 (0.337) | 0.939 (0.205) |
| CC* | 0.982 (0.71) | 0.984 (0.584) |
| Reflections used in refinement | 12071 (2243) | 55160 (3955) |
| Reflections used for R-free | 725 (135) | 1999 (143) |
| R-work | 0.2438 (0.2831) | 0.2572 (0.3945) |
| R-free | 0.2755 (0.2974) | 0.2937 (0.3878) |
| Number of non-hydrogen atoms | 6298 | 13206 |
| macromolecules | 6293 | 13067 |
| ligands | 5 | 0 |
| solvent | 0 | 139 |
| Protein residues | 807 | 1728 |
| RMS(bonds) | 0.002 | 0.003 |
| RMS(angles) | 0.45 | 0.68 |
| Ramachandran favored (%) | 93.22 | 92.78 |
| Ramachandran allowed (%) | 5.90 | 6.51 |
| Ramachandran outliers (%) | 0.88 | 0.70 |
| Rotamer outliers (%) | 0.00 | 2.79 |
| Clashscore | 6.42 | 10.03 |
| Average B-factor | 76.98 | 69.23 |
| macromolecules | 76.97 | 69.47 |
| solvent/ligands | 82.77 | 47.05 |

**Table 3.** Contracts of AF6-88.5 to H2-K^b^.

| H2-K<sup>b</sup> | AF6-H CDR | Distance (Å) | H2-K<sup>b</sup> | AF6-L CDR | Distance (Å) |
| --- | --- | --- | --- | --- | --- |
| R111 | D62 2 | 2.87 | R111 | D1 | 2.96 |
| Y113 | K65 2 | 3.14 | D212 | W94 3 | 3.43 |
| E128 | D62 2 | 2.80 | Q218 | K49 2 | 2.99 |
| E128 | K65 2 | 3.58 | G221 | S56 2 | 3.53 |
| T134 | G66 2 | 3.52 | E222 | S56 2 | 3.22 |
| I213 | G101 3 | 3.54 | E223 | K49 2 | 3.77 |
| T214 | T100 3 | 3.33 | E223 | I55 2 | 3.51 |
| T214 | G101 3 | 3.02 | Y262 | K49 2 | 3.72 |
| T214 | S102 3 | 3.09 | Y262 | Y50 2 | 3.15 |
| L215 | T100 3 | 3.63 | Y262 | E53 2 | 2.79 |
| T216 | T100 3 | 3.21 | P269 | E53 2 | 3.38 |
| T216 | W103 3 | 3.82 |  |  |  |
| E223 | W103 3 | 2.85 |  |  |  |
| E223 | A105 3 | 3.75 |  |  |  |
| I225 | F27 1 | 3.14 |  |  |  |
| I225 | F32 1 | 3.56 |  |  |  |
| I225 | A105 3 | 3.85 |  |  |  |
| I225 | Y106 3 | 3.52 |  |  |  |
| Q226 | D1 | 2.96 |  |  |  |
| Q226 | V2 | 3.89 |  |  |  |
| Q226 | G26 1 | 3.15 |  |  |  |
| M228 | T28 1 | 3.76 |  |  |  |
| M228 | F32 1 | 3.35 |  |  |  |
| E229 | S31 1 | 3.56 |  |  |  |
| L230 | S31 1 | 2.86 |  |  |  |
| L230 | F32 1 | 3.86 |  |  |  |
| E232 | S52 2 | 3.74 |  |  |  |
| E232 | S53 2 | 3.58 |  |  |  |
| E232 | G54 2 | 3.09 |  |  |  |
| E232 | N56 2 | 2.91 |  |  |  |
| Y262 | W103 3 | 3.37 |  |  |  |

| β<sub>2m</sub> | AF6-H CDR | Distance (Å) |
| --- | --- | --- |
| T28 | N56 2 | 3.46 |
| Q29 | G54 2 | 3.09 |
| Q29 | S55 2 | 2.73 |
| Q29 | N56 2 | 3.26 |
| S57 | S55 2 | 3.34 |
| S57 | N56 2 | 3.34 |
| K58 | I58 2 | 2.84 |
| K58 | Y59 2 | 3.79 |
| K58 | Y60 2 | 3.67 |
| K58 | K65 2 | 2.92 |
| D59 | S55 2 | 2.69 |
| D59 | I58 2 | 3.29 |

**Interface statistics**
| Chains | No. Interface Residues | Interface area ( Å<sup>2</sup>) | No. salt bridges | No. H bonds | No. contacts |
| --- | --- | --- | --- | --- | --- |
| A:H | 16:20 | 943:880 | 1 | 8 | 109 |
| A:L | 8:8 | 357:358 | 1 | 3 | 39 |
| B:H | 5:7 | 263:245 | - | 2 | 31 |

### X-ray Structures of the AF6-88.5/H-2K^b^ complex and AF6-88.5 alone

The X-ray structure of the Fab AF6-88.5/H-2K^b^ complex, even at at a lower overall resolution than the cryo-EM structure, is essentially identical to the cryo-EM structure. Data collection and refinement statistics are provided in Table 2, indicating that the complex was determined in space group P2_1_2_1_2_1_ to a resolution of 3.44 Å (Fig. 1*E*), and the unliganded Fab fragment of AF6-88.5 in space group P2_1_ to a resolution of 2.6 Å (Fig. 1*F*). The AF6-88.5/H2-K^b^ complex has five chains (Fab H and L, H2-K^b^, mβ_2_m, and peptide) in the AU (PDB-ID: 9Q0K), clearly establishing the same region of contact of AF6-88.5 with H-2K^b^ as seen in the model derived from the cryo-EM data (PDB-ID: 36FZ). The region of the α2-1 helix seen in the X-ray map reveals good density consistent with the model. Contacts as determined from the X-ray structure are the same as those determined from the cryo-EM model. AF6-88.5 alone crystallized with four HL complexes in the AU (PDB-ID: 9Q0L).

Examination of the structure of the complex in light of the allele and species specificities exhibited by the mAb indicates that the differences between mouse and human β_2_m as well as polymorphic differences in the contacts to the α2 domain (Q114) and to the large interface with the α3 domain (positions I225, D227 and Y262) are critical to the interaction (see Fig. 2*C* and Fig. 4).

**Figure 4.**
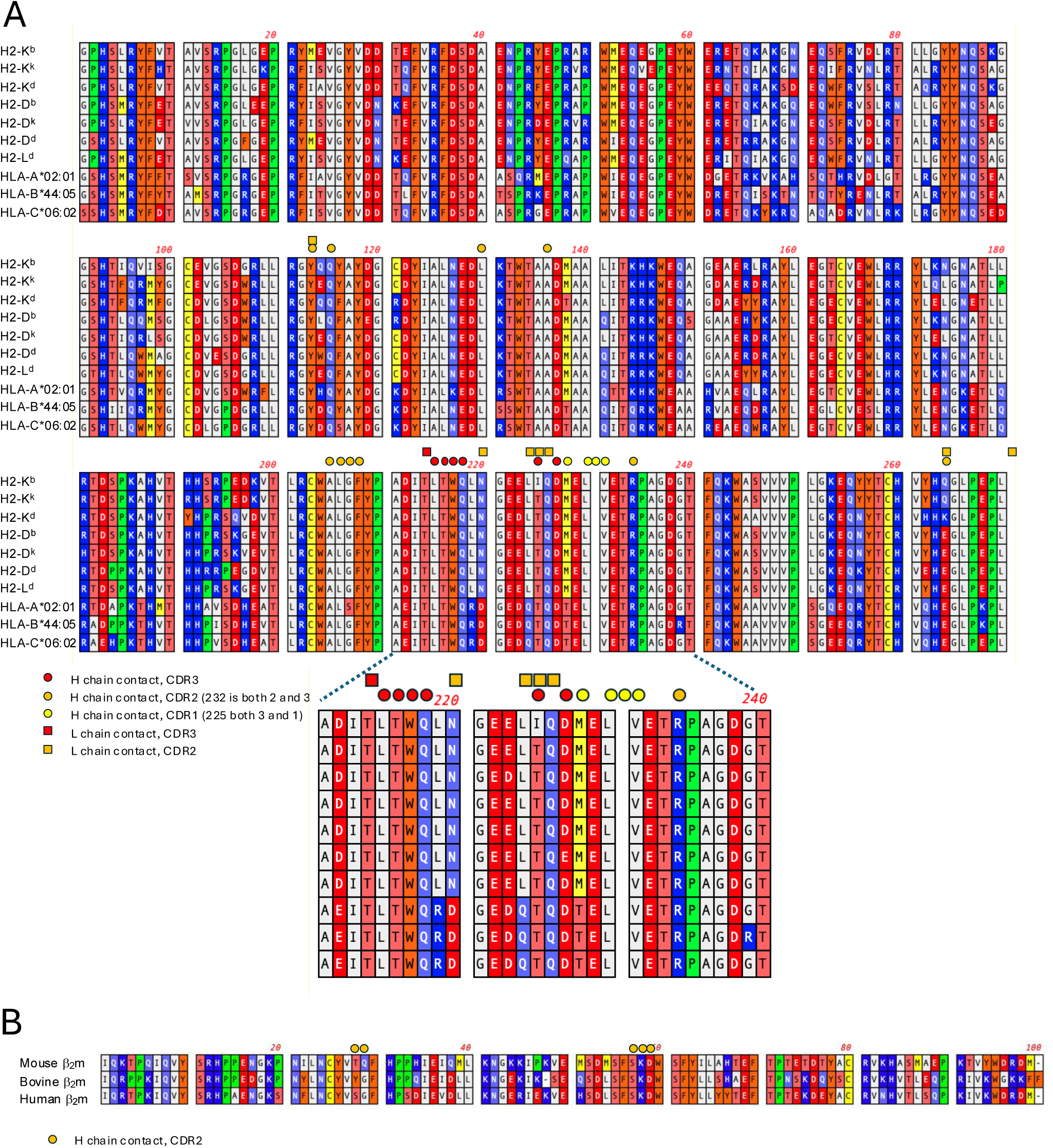
Amino acid sequence comparison reveals critical contacts of AF6-88.5to H2-K^b^. *A,* Amino acid sequences of the mature H2 or HLA heavy chain extracellular domains, or *B,* of the indicated species of β_2_m light chains were aligned with ClustalW^40^. Amino acid residues that contact AF6-88.5 are indicated. According to a more complete set of mouse MHC-I sequences^41^, I225 is unique to H2-K^b^, while all others, haplotypes f, k, p, r, s, v, w28, and alleles D^b^, D^dx^, D^f^, D^k^, D^p^, D^q^, D^r^, D^s^, L^d^, L^q^, D/Lv1, D/Lv2, D/Lv3 have T at that position.

Comparison of the unliganded AF6-88.5 X-ray structure (PDB-ID: 9Q0K) with its complexed counterpart (PDB-ID: 9Q0L) reveals remarkable similarity, with an RMSD of 0.680 Å for the AF6-88.5 H chain and of 0.827 Å for the AF6-88 L chain. Although there are slight changes observed for the AF6-88.5 H and L chain contacting residues in the CDR loops, the greatest change is in the adjustment of W103 of the AF6-88.5 H chain which rotates about the CB to CG bond repositioning the NE1 2.5 Å to form an H bond with OE2 of E223 of H2-K^b^. This rearrangement moves the CZ3 atom of W103 as much as 6.1 Å (see Fig. 3*A*). The unique I225 that defines H2- K^b^ specificity (see below) nestles into a hydrophobic patch lined by F27, F32, A105 and A106 of the AF6-88.5 H chain (Fig. 3*B*). The contacts of AF6-88.5 with the MHC-I light chain, β_2_m, are also significant (Fig. 3*C*), since H2-K^b^ complexed with human β_2_m revealed a low apparent affinity and higher dissociation rate (Fig. 1*A*). Detailed views of the superposed X-ray (PDB-ID: 9Q0K) and cryo-EM models (PDB-ID: 36FZ) are illustrated in Fig. 3*D*, with an overall RMSD of 1.39Å.

In comparing the density maps derived from the cryo-EM and X-ray data, we observed that the cryo-EM map lacked some density on the surface in the region of the α2-1 helix. Nevertheless, the density in the contact region at the interface between the mAb and H2-K^b^ is well defined (Fig. 3*E* to 3*I*) where the local resolution is highest (Fig. S1F).

### Sequence comparison and mutational analysis

To understand the structural basis for the specificity of AF6-88.5 for H2-K^b^/mβ_2_m complexes, we compared the amino acid sequences of selected murine H2 and human HLA molecules and murine, human and bovine β_2_m chains (see Fig. 4, *A* and *B*). Consideration of the MHC-I heavy chain residues that are contacted by AF6-88.5 focuses attention to I225 of H2K^b^. Comparison of β_2_m sequences (taking note of the lack of reactivity of AF6-88.5 with H2-K^b^ bound to bovine (22) or human (16, 22) β_2_m) directs us to the critical roles of residues S28 and G29. I225 of H2-K^b^ is unique among mouse and human MHC-I molecules. Residues T28 and Q29 of β_2_m are contact residues to the AF6-88.5 H chain that differ between human, bovine, and mouse.

To address the hypothesis that the unique polymorphism of I225 in H2-K^b^ and the T28/Q29 species differences of mβ_2_m as compared with hβ_2_m are major contributors to the specificity of AF6-88.5, we generated several site-directed mutations of H2-K^b^ and H2-D^b^, as well as of mβ_2_m and hβ_2_m. Proteins were expressed, assembled with appropriate H2-K^b^ or H2-D^b^ specific peptides, and subjected to quantitative binding analysis to AF6-88.5 by SPR. As shown in Fig. 5, H2- K^b^/mβ_2_m binds AF6-88.5 with a *K*_D_ of ∼0.4 μM while H-2D^b^/mβ_2_m, consistent with qualitative analysis (Fig. 1*A* and (16)), shows no detectable binding. Mutation of the single focal I225 residue of H2-K^b^ to T abolished binding, and conversely, mutation of the weakly binding H2-D^b^ T225 to I improved its affinity to about 3.1 μM. Mutation of mβ2m T28/Q29 to its human counterpart S28/G29 reduced the affinity to an undetectable level and, conversely, mutation of hβ_2_m S28/G29 to its mouse counterpart T28/Q29 improved the apparent binding, but under these conditions this could not be accurately quantified.

**Figure 5.**
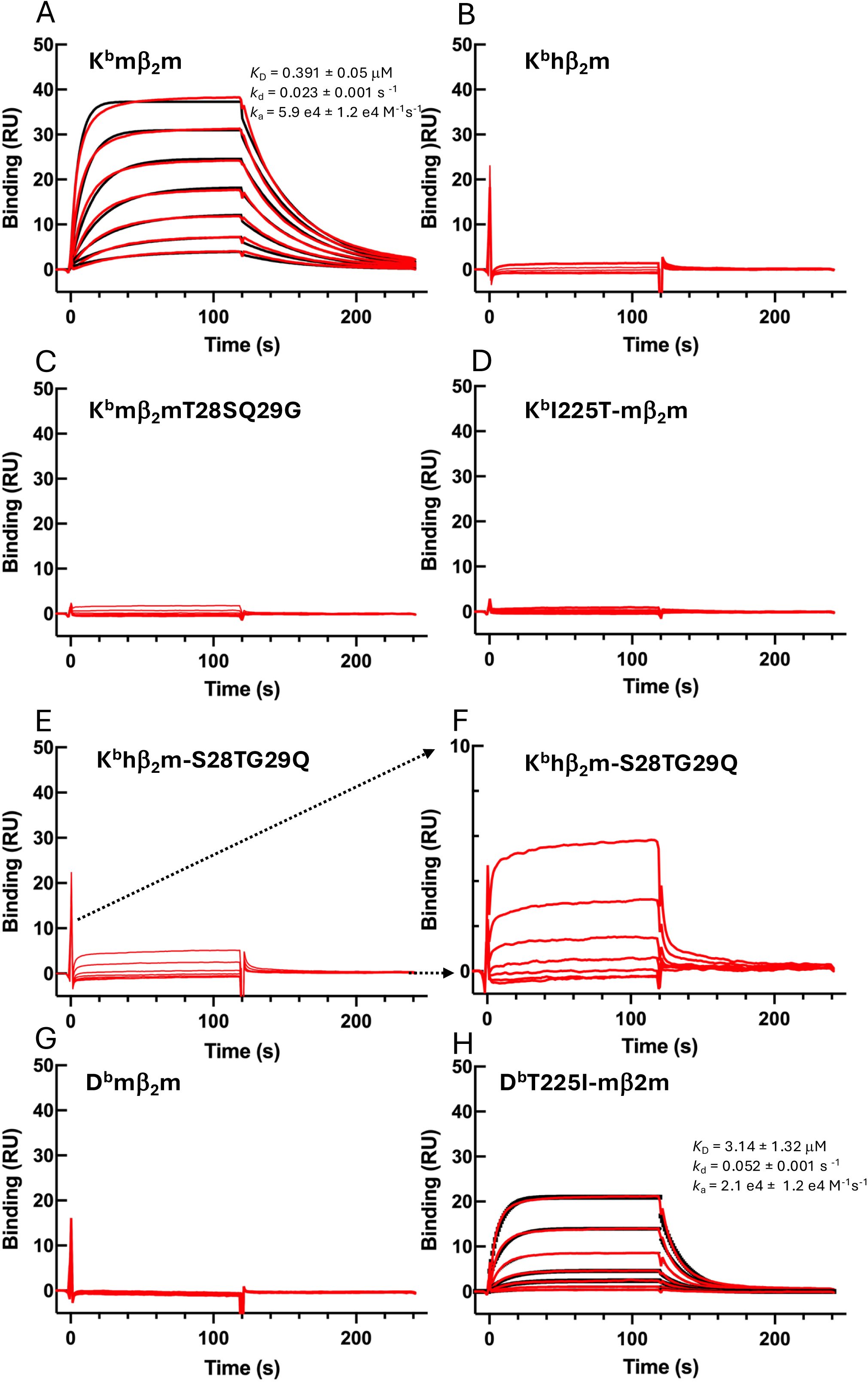
Binding of site-directed mutants verifies role of I225 of H2-K^b^ and of T28 and Q29 of β_2_m in specificity. SPR binding studies of each of the indicated H2/β_2_m/peptide complexes to an AF6-88.5 coupled surface were performed as described in **Experimental procedures.** Complexes were: *A,* H2-K^b^/mβ_2_m/p; *B,* H2-K^b^/humanβ_2_m/p; *C*, H2-K^b^/mβ_2_m T28S, Q29G mutant/p; *D*, H2-K^b^ I225T mutant/mβ_2_m/p; *E,* H2-K^b^/hb_2_m S28T, G29Q mutant/p; *F,* expanded plot of *E* showing concentration dependent binding; *F*, H2-D^b^/mβ_2_m/p; *G,* H2-D^b^ T225I mutant/mβ_2_m/p. Calculations of *k*_a_, *k*_d_, and *K*_D_ obtained from single site kinetics are shown for *A* and *H*.

## Discussion

The ability of antibodies to discriminate subtle amino acid sequence differences among protein antigens has been known for many decades and is the basis of classical tissue typing (24) and analysis of immunoglobulin idiotypes and allotypes (25–27). Monoclonal antibodies directed against MHC antigens are invaluable not only for their immunological applications to identify structural similarities and differences but also for their functional uses in blocking T cell and NK receptor interactions and co-receptor inhibitory and activating effects. Facility in determining molecular and atomic resolution details of mAb/protein antigen complexes fosters a deeper understanding of the general rules of antibody recognition and the broader themes that define protein antigenicity. Because of the huge database of MHC-I (particularly human HLA-A, -B, - C) sequences (6), the accumulation of detailed information concerning specific anti-MHC mAbs may not only help to define epitopic sites but also to refine the ability to computationally predict such sites from mAb amino acid sequence alone. Conversely, expanding our knowledge of the details of highly discriminatory mAb/protein interactions may contribute to the future potential to computationally design focused binding molecules of mAb, nanobody, miniprotein, or peptide composition. Here, we studied in biochemical and structural detail one mAb, AF6-88.5, with previously reported specificity for the murine MHC-I molecule, H2-K^b^. Binding studies using purified mAb and recombinant MHC-I were consistent with previously reported specificity analysis and confirmed the known preference for H2-K^b^ bound to murine β_2_m and peptide. The cryo-EM and by X-ray crystallography structures revealed the same footprint, supporting the view that the two distinct modes of analysis, one based on solution and one based on crystalline structure, reveal equivalent structures.

The two independently determined AF6-88.5 Fab/H2-K^b^ structures provide complementary views of different parts of the complex. Of note, the carboxyl end of the α1 helix (residues 73 to 91) and the amino end of the α2 domain (the α2-1 helix, residues 138 to 151) fit into rather weak coulombic density in the cryo-EM map. In general, although the cryo-EM map overall is at a higher resolution than the X-ray electron density map, the local resolution in the cryo-EM map resolution is variable, with highest resolution at the mAb/H2-K^b^ interface and lower resolution in the α1 and α2 domain residues furthest from the interface. This is illustrated by the *Q*-score metric which evaluates the fit of various regions of the model to the map (28, 29)(Fig. S2). Comparison of the final model with the cryo-EM map overall gives a *Q*-score value of 0.35, but chain by chain and domain by domain comparison reveals that different regions of the complex structure are resolved with greater or lesser precision. Specifically, the mAb H and L chain model better (*Q-*scores 0.48 and 0.45 respectively) than the H2-K^b^ heavy chain (Fig. S2*A*) (*Q-*score 0.22) or β_2_m (Fig. S2*B*) (*Q*-score 0.29). Also, the distinct domains of H2-K^b^ progress from rather poor *Q*-score for the α1 (0.09) and α2 (0.12) domains (on the surface of cryo-EM map) as compared to α3 (0.44) which interfaces with the mAb (mass center). These differences are seen in the per residue *Q*-scores plotted in Fig. S2*B*. The plots of Q-scores also associate with the “local resolution plot” presentation at Fig. S1*F*.

Comparison of the bound mAb with its unliganded counterpart showed minor but detectable conformational adjustments of the mAb to accommodate the MHC-I molecule. This finding indicates that the intrinsic dynamics of the mAb could fine tune its binding interface to recognize antigen with higher affinity rather than relying on a rigid lock and key interaction. Finally, key amino acid contact residues of H2-K^b^ and mβ_2_m, when mutated could abolish binding, while reactivity could be recovered by judiciously engineered mutations in the non-binding H2- D^b^ and hβ_2_m. The site here identified structurally refines the work of Kuhns and Pease (16) who found little effect on AF6-88.5 binding of a set of H2-K^b^ mutant mice. However, their determinations that G162D decreased binding and that N174K increased binding to AF6-88.5 are not easily interpretable in light of the structures we present here.

The site on H-2K^b^ recognized by AF6-88.5 largely overlaps the site utilized on this and other MHC-I molecules by some human and murine receptors, some mAbs, and by a viral immunoevasin. The AF6-88.5 footprint on H2-K^b^ overlaps the binding site of the coreceptor, CD8 (both CD8αα and CD8αβ (PDB 1BQH (30) and 3DMM (31) respectively), accounting for its ability to discriminate high from low affinity TCR interactions. In addition, the AF6-88.5 site sterically blocks the NK inhibitory receptor Ly49C binding site (PDB 3C8K (32)). Also, the cowpox immunoevasin CPXV203 binds in a site overlapping AF6-88.5 (33). These authors also pointed out that H2-K^b^ M228T (a CDRH1 contact site and a polymorphism that distinguishes murine from human MHCI) obliterates binding to AF6-88.5. The analogous α3 domain region of H2-D^b^ and H2-L^d^ is seen by mAb 28-14-8 (PDB 8TQA (8)). Looking at the homologous HLA MHC-I molecules, the AF6-88.5 footprint is common to that of the inhibitory receptors LILRB1 (PDB 6EWA(34) and 5KNM(35)), and the anti-HLA mAbs W6/32 (PDB 7T0L(36)) and DX17 (PDB 8TQ5 (10)).

It is intriguing that a notable biological effect of AF6-88.5 is that it synergizes with the mAb 34-1-2S to cause lung damage in a murine model of transfusion-related acute lung injury (TRALI). TRALI is antibody and complement dependent and is not observed in mice receiving either antibody alone (21). Since the 34-1-2S epitope has been localized to the α1 domain of H2- K^d^ (37) and the complement dependent activation of TRALI has been modeled based on the formation of hexameric Ig clusters resulting from 34-1-2S (38), we speculate that the synergistic effect of AF6-88.5 is due to topological arrangements of cell surface H2-K^b^ molecules that permit binding of 34-1-2S orienting its Fc to be available for interaction with the C1q hexamer. We visualize the binding of AF6-88.5 to surface H2-K^b^ molecules in a manner that would dimerize them via the α2α3 binding site, orienting the α1α2 domain surface away from the cell membrane. Thus, an antibody such as 34-1-2S that binds the α1 domain of H2-K^b^ (mapped for H2-K^d^ (39) and H2-D^d^ (40)) at low affinity (21) would then bind to orient its Fc portion for access by C1q, initiating a complement cascade. Structural analysis of this AF6-88.5/MHC complex not only defines the epitope, it offers an explanation for the unique role that AF6-88.5 plays in the C57BL/6 TRALI disease model.

## Experimental procedures

### Antibodies, purification, and preparation of Fab

AF6-88-5.3 cells (HTB 158) were obtained from the American Type Culture Collection (ATCC), propagated in DMEM containing 10% fetal bovine serum, 100 mM glutamine, non-essential amino acids at 37 °C in a humidified atmosphere. Supernatants were harvested and secreted antibody was purified on Protein A-Sepharose as described previously. (AF6-88.5.3 is a IgG2a murine monoclonal antibody obtained in a fusion of SP2/0 myeloma cells with spleen cells from BALB/c mice immunized with spleen cells from C57BL/10J mice that is considered to react only with H2-K^b^ and not with H2 of d, f, g, k, p, q, r, s, u, or v haplotypes (20)). Fab fragments were prepared from the purified IgG as described (41). Purified AF6-88.5 Fab was incubated with Kb+SIINFEKL+mβ_2_m at RT for ∼4 hours, in a molar ratio of 2 Fab:1MHC. The complex was purified on Superdex^TM^ 200 increase column, 10/300 (Cytiva) equilibrated in 1XPBS. Peak fractions were pooled and concentrated using Millipore 10 kDa spin concentrators. SDS-PAGE analysis confirmed presence of both Fab and MHC in peak.

### Cryo-EM sample preparation and data collection

Freshly purified anti-mouse antibodies (“Fab AF6-88”) were incubated in a 1:1 molar ratio with bacterially expressed and refolded soluble peptide/H2-K^b^/hβ_2_m complexes prepared as described (42). The complexes were purified by size exclusion chromatography (SEC) and used at a concentration of 0.7-1.4 mg/ml for sample preparation. Samples were applied onto the grids (Quantifoil® Ultra Au Foil R1.2 μm/1.3 μm space 300 mesh (EMS.com)), which had been glow discharged for 60 seconds, blotted for 3 seconds, and plunged into liquid ethane with a Vitrobot Mark IV (Thermo-Fisher) at 4 °C and 95% humidity. Cryo-EM data on the selected regions with ideal ice thickness were collected on a Titan Krios 300-keV microscope. Images were acquired automatically with SerialEM (43) on a BioQuantum-K3 detector (Gatan) in super-resolution mode at 105kx nominal magnification (0.83 Å/binned pixels) and a nominal defocus range from −0.7 to −2.0 μm. An exposure time of 0.05s per frame was recorded, with a total exposure of about 54.2 electrons/Å^2^. The data set was collected: AF6-88Fab+H2-K^b^ with 4,356 movies.

### Image processing, map resolution improvement, and model refinements

All image processing, 2D class, 3D reconstruction, and map refinements were performed using cryoSPARC v4.7.3 (23, 44, 45), and updated by using CryoSPARC^TM^ v5.0.3. Following “Patch Motion Correction,” “Patch CTF Estimation,” and “Curate Exposures,” outliers of defocus range, defective micrographs, and low-resolution estimation of the CTF fit (>5 Å) were discarded. The “Blob Picker” was initially used with a particle diameter of 110 Å for picking particles. The box size used for 2D classification and following was 240 pixels. The initial “Blob Picker” resulted in only a few 2D classes with suitable particles. Subsequently, we used these initial 2D classes as templates for “Template Picker” and, following standard protocols (“Ab-Initio Reconstruction” and “Non-uniform Refinement”) with several iterations, obtained a map resolution of about 4.0 Å for these antibody complexes. We used a multiple Low-Pass Filter (MLPF) protocol (Jiang et al, 2026) for map resolution improvement in cryoSPARC(23) as shown in Extended Data Fig.2. The extracted particles are initially 2,326,183 particles, and final 359,876 particles were used for the final refined map (Fig.2d). The map resolution was improved to 3.09 Å with FSC at 0.143 (see Fig. S1 and Table 3).

We used the X-ray crystal structure model (PDB: 9Q0K) to dock and manually fit the cryo-EM maps of AF6-88Fab+H2-K^b^by ChimeraX(46), then we used Real-Space Refinement in Phenix (47), which includes rigid-body refinement and with the secondary structure restraints. The MHC-I rigid-body domains consist of α1α2+peptide, α3, and β_2_m domains, and Fab consists of four rigid-body domains (V_L_, C_k_, V_H_, C_H1_). Simulated annealing (SA) at the initial step, local grid search, and ADP refinement were included. The final refined model compared with the map densities has an overall CC (Correlation Coefficient) of 0.75/0.73/0.62 (mask/volume/peaks) for AF6-88Fab+H2-K^b^. We also calculated *Q*-score of individual residues (29) for validation. Cryo-EM Data processing, refinement statistics, and model validation are listed in Table 1.

### Crystallization and Data Collection, and Refinement

Crystallization conditions were identified by screening hanging drops at 18 °C. Crystals of AF6- 88+H2-K^b^ were obtained in 18% PEG 3350, 0.1M Na citrate, pH 5.6, 0.2M LiSO_4_. The crystals appeared in grain-like and pin-like/needle-like shapes. X-ray diffraction screening found that the grain-like crystals represented H2-K^b^ alone whereas the pin-like/needle-like crystals were the complex of AF6-88 Fab and H2-K^b^. However, the resolution for the complex initially was only at 4.0 Å. The crystals for the complex were optimized, improving the data resolution to 3.44 Å. The crystals of AF6-88.5 alone were obtained in 12% PEG 4000, 0.1M Tris, pH 6.8, 10% glycerol. The crystals appeared as 2D plates with multiple layers and diffracted to 2.40 Å resolution. However, as the data quality in the high-resolution range was poor, a resolution cutoff of 2.60 Å was applied which resulted in a lower R_free_ value.

Crystals were cryoprotected in mother liquor containing 10% ethylene glycol and flash frozen in liquid nitrogen. Diffraction data were collected (at wavelength 1.000 Å, in N_2_ stream at ∼ 100 K) at Southeast Regional Collaborative Access Team (SER-CAT) beamline 22ID at the Advanced Photon Source, Argonne National Laboratory, and processed with XDS (48). The complex structure was solved by molecular replacement with Phaser (49). The search model of H2-K^b^ is taken from PDB 1VAC, and the AF6-88.5 model was obtained by the Alphafold3(50) prediction with the CDR loops trimmed off. Then the CDR loops were manually built according to the electron density. These molecular replacement models were subjected to several rounds of refinement with Phenix (51) interspersed with manual building in Coot (52). The AF6-88.5 Fab sequences were determined by RT-PCR and sequencing of the hybridoma cDNA. R_work_/R_free_ (%) values for the final refined model of the complex of AF6-88.5 Fab and H2-K^b^ is of 24.4/27.6, and AF6-88.5 Fab alone is of 25.7/29.4. The structure of AF6-88.5 Fab alone has four copies of the molecule in the asymmetric unit of the P2_1_ space group. Data collection and refinement statistics are summarized in Table 2. Figures were generated with PyMOL (53) and ChimeraX (46), and sequence comparisons were performed in MacVector v. 18.8.3 (32) using ClustalW (54). Additional mouse H2 sequences were taken from Pullen et al (55).

## Data availability

Cryo-EM map and model of AF6-88.5Fab+H2-K^b^ have been deposited at the Electron Microscopy Data Bank (EMDB) and Protein Data Bank (PDB) under accession codes 77500 and 36FZ respectively. The X-ray data and models of the AF6-88.5Fab+H2-K^b^complex and the unliganded AF6-88.5 Fab are deposited at the PDB under accession codes 9Q0K and 9Q0L respectively.

## Supporting information

This article contains supporting information.

## Acknowledgements

We appreciate discussions with Shikha Sharma and Martin Meier-Schellersheim. This study used the Office of Cyber Infrastructure and Computational Biology (OCICB) High Performance Computing (HPC) cluster at the National Institute of Allergy and Infectious Diseases (NIAID), Bethesda, MD. X-ray data were collected at Southeast Regional Collaborative Access Team (SER-CAT) 22-ID beamline at the Advanced Photon Source, Argonne National Laboratory. SER-CAT is supported by its member institutions (www.ser-cat.org/members.html) and equipment grants (S10_RR25528 and S10_RR028976) from the National Institutes of Health. Use of the Advanced Photon Source was supported by the U. S. Department of Energy, Office of Science, Office of Basic Energy Sciences, under Contract No. W-31-109-Eng-38. The Electron Microscopy Resource is supported by the National Cancer Institute and NIH Intramural Research Program Cryo-EM Consortium (NICE).

## Funding and additional information

This research was supported by the Intramural Research Program of the National Institutes of Health (NIH). The contributions of the NIH author(s) were made as part of their official duties as NIH federal employees, are in compliance with agency policy requirements, and are considered Works of the United States Government. However, the findings and conclusions presented in this paper are those of the author(s) and do not necessarily reflect the views of the NIH or the U.S. Department of Health and Human Services.

## Supporting Information Figure

**Figure S1.**
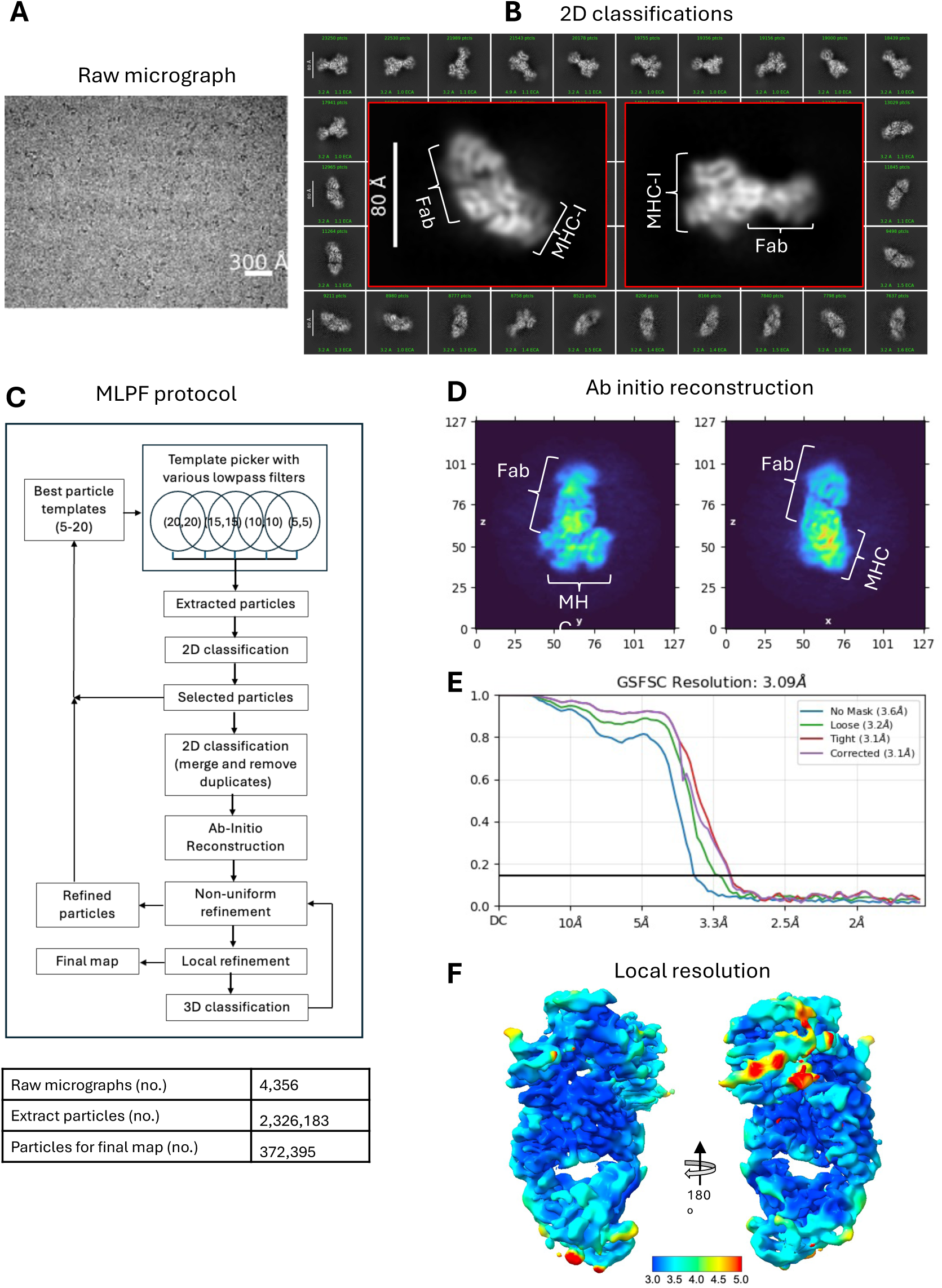
Data processing and evaluation of cryo-EM structure of AF6-88.5+H2-K^b^. *A,* Example of raw micrograph (data set: 4,356 movies). *B,* 2D classification (50x), insertions with the two enlarged 2D class average that clearly show AF6-88 in contact with H2-K^b^. *C,* MLPF protocol used in the particle picker with CryoSPARC^TM^ (23). *D, Ab initio* reconstruction with two typical orientations. *E,* FSC evaluation of map resolution. *F,* Local resolution representation.

**Figure S2.**
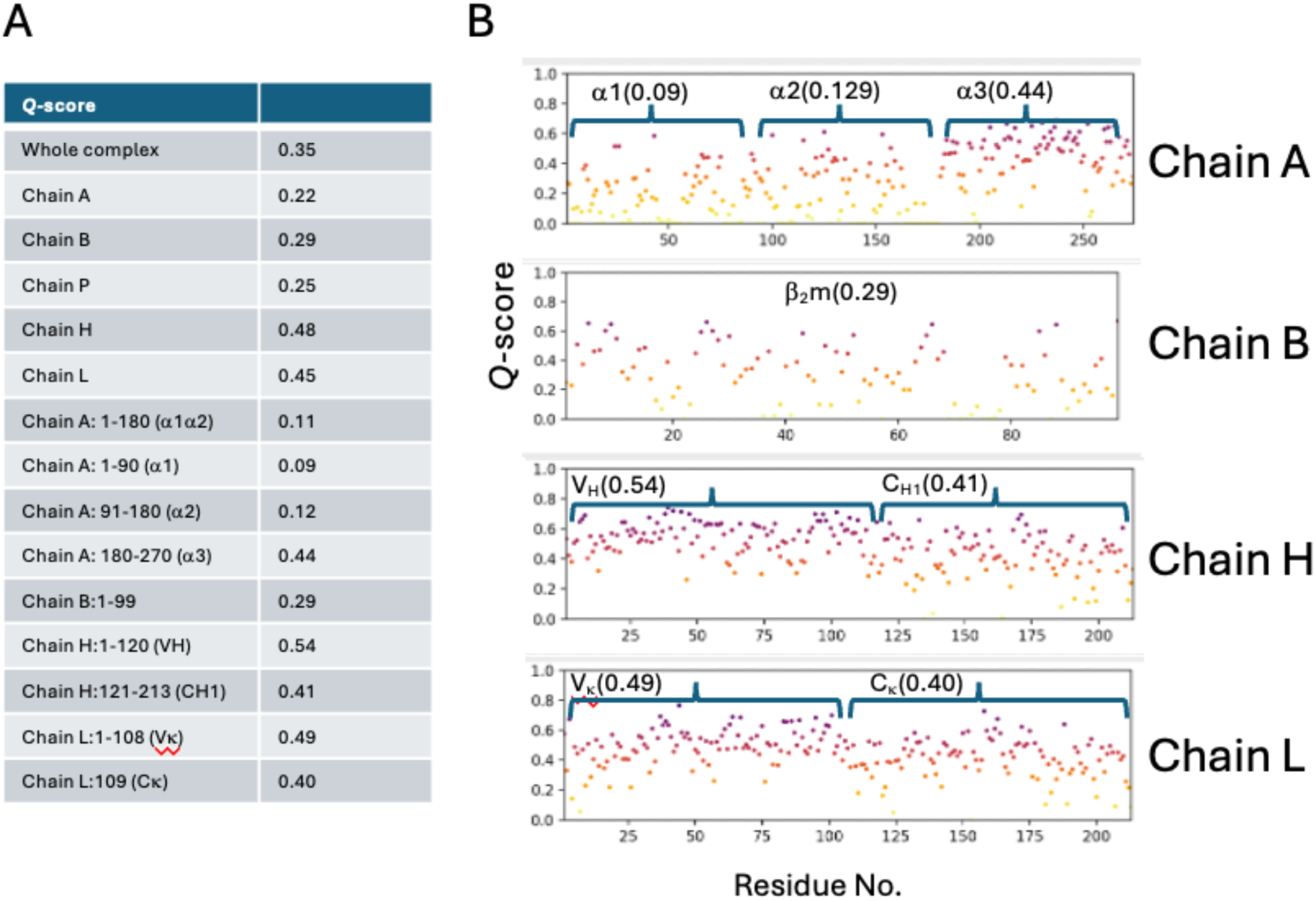
*Q*-scores comparing map and model show regions of greater and lesser fidelity of fit. *A,* Q-scores comparing the model and the cryo-EM map were calculated in Chimerax^TM^(23) for the indicated chains as well as the indicated domains of the proteins. *B*, per residue *Q*-scores are shown, indicating the domains of the proteins and the average *Q*-score for the regions.

